# Image-based profiling can discriminate the effects of inhibitors on signaling pathways under differential ligand stimulation

**DOI:** 10.1101/190637

**Authors:** Kenji Tanabe, Yuji Henmi, Masanobu Satake

**Affiliations:** Medical Research Institute, Tokyo Women’s Medical University, Tokyo, Japan; School of Nursing, Sendai Akamon College, Sendai, Japan

**Author notes:** Correspondence should be addressed to Kenji Tanabe. Medical Research Institute, Tokyo Women’s Medical University, 8-1, Kawada-cho, Shinjuku-ku, Tokyo 162-8666, Japan.

**Keywords:** High contents screening, signal transduction, Cell-based assays, Phenotypic drug discovery, Cancer and cancer drugs

## Abstract

A major advantage of image-based phenotypic profiling of compounds is that numerous image features can be sampled and quantitatively evaluated in an unbiased way. However, since this assay is a discovery-oriented screening, it is difficult to determine the optimal experimental set-up in advance. In this study, we examined whether variable cellular stimulation affects the efficacy of image-based profiling of compounds. Seven different EGF receptor ligands were used, and the expression of EGF receptor signaling molecules was monitored at various time points. Significant quantitative differences in image features were detected among the differentially treated samples. Next, 14 different compounds that affect EGF receptor signaling were profiled. Nearly half of the compounds were classified into distinct clusters, irrespective of differential ligand stimulation. The results suggest that image-based phenotypic profiling is quite robust in its ability to predict compound interaction with its target. Although this method will have to be validated in other experimental systems, the robustness of image-based compound profiling demonstrated in this work provides a valid basis for further study and its extended application.

## Introduction

In recent years, a main stream of drug discovery has been a target-based screening approach that aims to determine whether the function of a given target molecule is affected by a compound^1,2^. However, numerous candidate compounds selected are often abandoned early in the screening process. One reason for this may be that the target-based approach focuses solely on one target molecule and does not take into account the entire spectrum of biological processes^3,4^. Thus, a phenotype-based drug discovery approach that was popular many years ago has once again gained interest^5^, now integrated with new techniques, such as automated sampling of microscopic images combined with quantitative and statistical image analysis^6–8^. Since many image features can be obtained and analyzed, this type of phenotypic profiling is also called high content screening (HCS)^9^. These days, HCS is being applied to many facets of drug screening^6,10,11^.

Most of the recently reported image-based phenotypic profiling does not constitute bona fide HCS but rather “low content, high throughput” screening. For example, only a single image feature (e.g., cytoplasmic to nuclear translocation or neurite extension) has been selected and used for drug screening^12^. Because of its simplicity, this “low content, high throughput” screening has been successful in many research fields but does not make use of many, possibly useful image features. A major advantage of authentic phenotypic profiling/HCS is that many image features obtained from single cells can be quantitatively evaluated in an unbiased way.

There are several potential workflow hurdles in carrying out HCS^12^. The use of the Z’-factor, a criterion for measuring the separation between the positive and negative controls, is an example of these hurdles. For a single content screening, it would be rather easy to optimize experimental conditions by monitoring the Z’-factor, whereas, for HCS, it may not be immediately clear if one can achieve optimal experimental conditions by tuning a Z’-factor. Because HCS handles numerous image features and is a discovery-oriented screening, it is difficult to determine the optimal experimental set-up in advance. Theoretically, it would be ideal to perform HCS under all possible experimental conditions, but such an approach would be costly and unrealistic. If one can determine ahead of time how differential experimental conditions affect image-based phenotypic profiling of drugs, authentic HCS would constitute a worth-while and convenient approach.

Thus, the purpose of this study was to evaluate whether variant cellular stimulation affects the image-based profiling of different compounds. We chose several different EGF receptor (EGFR)^13–16^ ligands and monitored the expression of EGFR signaling molecules^17^. First, images of cells were compared by microscopy. Second, numerous image features were automatically extracted and analyzed. Significant qualitative differences in the images as well as quantitative differences in the image features were detected among the samples treated with the different ligands. Finally, when compounds that are known to affect EGFR signaling were profiled, the compounds were classified similarly irrespective of differential ligand stimulation. This result indicates that image-based phenotypic profiling/HCS is considerably robust in classifying inhibitors under differential cellular stimulation conditions.

## Materials and methods

### Cells and reagents

A549 GFP-EGFR cells that express GFP-tagged endogenous EGFR protein were purchased from Sigma (CLL1141, St. Louis, MO) and maintained at 37°C in DMEM containing 10% FBS and antibiotic-antimycotic solution (A5955, Sigma). Recombinant human EGF, HB-EGF, and transferrin were purchased from R&D Systems (Minneapolis, MN). Recombinant human ARG, BTC, EPGN, and ERG were from PeproTech (Rocky Hill, NJ). Human TGFα was from Cell Signaling Technologies (CST; Danvers, MA). Alexa Fluor 647-conjugated human transferrin and Hoechst 33342 were from Invitrogen (Carlsbad, CA). The following antibodies were used in this study: rabbit anti-phosphorylated ERK monoclonal antibody, rabbit anti-phosphorylated Akt monoclonal antibody (CST), murine anti-PtdIns(4,5)P_2_ monoclonal antibody (Echelon, Salt Lake City, UT), and rabbit anti-GST polyclonal antibody (Santa Cruz Biotechnology, Santa Cruz, CA). All Alexa Fluor-conjugated secondary antibodies were purchased from Invitrogen. Drugs were purchased as an EGFR inhibitor panel (Merck Millipore, Billerica, MA), and nocodazole was obtained from Sigma.

### Inhibitor treatment and ligand stimulation

Cells were seeded in 96-well plates (Edge-plates, Thermo Scientific, Waltham, MA), serum-starved for 6 h by replacing the medium with DMEM containing 0.1% BSA, and stimulated with EGFR ligands (100 nM AREG, 10 nM BTC, 100 ng/ml EGF, 250 ng/ml EPGN, 10 nM EREG, 10 nM HB-EGF, or 50 ng/ml TGFα). One hour prior to stimulation, the cells were treated with the following compounds at the indicated concentrations: Akt inhibitor VIII (200 nM)^18^, PD153035 (100 nM)^19^, CAS 879127-07-8 (10 μM)^20^, Et-18-OCH3 (5 μg/ml)^21^, JNK inhibitor II (1 μM)^22^, LY294002 (10 μM)^18^, PD98059 (50 μM)^23^, PD168393 (100 nM)^24^, PP2 (100 nM)^25^, rapamycin (10 nM)^26^, SB253080 (10 μM)^22^, AG490 (40 μM)^27^, ZM336372 (1 μM)^28^, and nocodazole (50 μM)^29^. Next, Alexa Fluor 647-conjugated transferrin and EGFR ligands were internalized for 5 min, followed by the addition of medium containing unlabeled human transferrin (R&D Systems) and the corresponding EGFR ligands. After the indicated time intervals (0, 5, 30, 60, and 180 min), the cells were fixed by adding an equal volume of methanol-free 4% paraformaldehyde (09154-85, Nacalai Tesque, Kyoto, Japan) for 15 min, washed three times with PBS, and subjected to immunofluorescence.

### Immunofluorescence

Fixed cells were prepared for immunofluorescence as previously described^17^. To stain phosphorylated ERK (pERK), fixed cells were incubated with ice-cold methanol for 10 min at −30°C, washed twice with PBS, and incubated in blocking buffer (1% BSA in PBS) containing 0.3% Triton X-100 (TX100) for 1 h at RT. To stain phosphorylated Akt (pAkt), fixed cells were incubated in blocking buffer containing 0.3% TX100. For staining phosphoinositides, fixed cells were permeabilized with 0.01% digitonin in PBS for 5 min at RT, washed once with PBS, and incubated in blocking buffer without TX100 for 1 h at RT. To label phosphatidylinositol 3-phosphate (PtdIns(3)P), recombinant GST-Hrs FYVE protein (prepared as described^17,30^) was applied after permeabilization with 0.01% digitonin for 1 h at RT, followed by two washes with PBS. The cells were incubated with the following primary antibodies diluted with block buffer for 1 h at RT: anti-pERK (1:1000), anti-pAkt (1:2000), anti-PtdIns(4,5)P_2_ (1:1000), or anti-GST (1:200). The cells were washed twice with PBS and then incubated for 1 h at RT with the appropriate secondary antibodies and Hoechst 33432. The cells were washed twice with PBS, fixed again with 2% paraformaldehyde for 5 min at RT, washed once with PBS, and stored at 4°C.

### Image acquisition and analysis

Stained cells were photographed using an automated fluorescence microscope (CellInsight, Thermo Scientific) with a 20× objective lens. Image analysis was performed as previously described^17^. Briefly, Hoechst and GFP signals were used to define the ‘nucleus’ and ‘cell’, respectively. Thirty-six fields of view were photographed for each well. Image analysis was performed using CellProfiler (Broad Institute, Cambridge, MA), and all images were processed for illumination correction. Using the nuclear (Hoechst) and GFP-EGFR signals, the cell outline (‘cell’) was identified by the ‘propagation’ implemented in CellProfiler. ‘Cytoplasm’, ‘perinuclear’, and ‘PM’ were defined from ‘cell’ and ‘nucleus’ as previously described^17^. These regions of interest (ROIs) were used to quantify each image feature. Some fluorescence signals were converted into objects to identify ‘endosomes’ using the Top-Hat filter implemented in the ‘enhance’ module of CellProfiler. Co-localization was also evaluated using these objects. The image features used in this study are listed in Supplementary Table S1.

### Data processing and statistical analysis

Image analysis data were stored on a local MySQL server and subjected to statistical analysis using MatLab (Mathworks, Natick, MA). To evaluate ligand stimulation, image features of each ligand were compared against unstimulated (0 min) cells using the modified two-sample Kolmogorov-Smirnov (KS) test and the standardized Z-scores were calculated^6,17^. Mean Z-scores were obtained for two independent experiments. When performing compound profiling in the ligand-stimulated cells, image features for each drug treatment were compared with the control (DMSO-treated) samples and processed as described above to obtain mean Z-scores. Next, the mean Z-scores were subjected to principal component analysis (PCA). All PCs were used for subsequent hierarchical clustering with the cosine distance and the average-linkage methods.

## Results

### Differential cellular responses induced by EGFR ligands can be recognized at the level of raw microscopic images

To stimulate the cells, seven different ligands that can bind to EGFR were employed. According to the literature, these ligands may be tentatively classified into two groups based on receptor-trafficking pathways^13^. One group, the members of which are termed degradative ligands, includes EGF, BTC, and HB-EGF. EGFR, when bound to a “degradative ligand”, is trafficked to an endosome-lysosome pathway and is eventually degraded. Another group, the members of which are termed recycling ligands, includes TGFα, AREG, EREG, and EPGN. EGFR, when bound to a “recycling ligand”, is incorporated into recycling endosomes and eventually recycled back to the plasma membrane. Furthermore, the degree of receptor phosphorylation and/or activation of downstream signals vary for the different ligands^14^. Thus, these different molecules are expected to induce distinct cellular responses through the same receptor.

First, the possibility of differentiating between cellular responses by evaluating microscopic images was assessed. For simplicity, each ligand was used at a fixed concentration (as described in the materials and methods section). EGF, which can direct EGFR toward both the degradation and recycling pathways, was employed at the concentration of 100 ng/ml^31^. A549 cells, harboring GFP-fused, endogenous EGFR, were serum-starved for 6 h, exposed to the ligand, and fixed after 5, 30, 60, or 180 min. Four major signal transducers, including pERK, pAkt, PtdIns(3)P, and phosphatidylinositol 4, 5-bisphosphate (PtdIns(4,5)P_2_), were visualized by immunofluorescence. The GFP-EGFR signal was used to visualize the trafficking of EGFR from the plasma membrane to the endosomes. To visualize recycling pathways, alexa647-conjugated human transferrin was employed.

Representative images showing GFP-EGFR and pAkt are presented in Fig. 1. When EGF, BTC, or HB-EGF was used for stimulation, endosome-like, high GFP-EGFR signal was clearly observed at 30 min. but not at 180 min. Considering that EGF-EGFR is generally transported to late endosomes about 30 min after binding^32^, the low GFP-EGFR signal at 180 min suggests that EGFR is degraded in the lysosomes. In parallel, high pAkt fluorescence signal was observed until 60 min and was lower at 180 min. On the other hand, stimulation by TGFα or AREG, the recycling ligands, resulted in a different signal pattern. A weak but significant, endosome-like GFP-EGFR signal was observed at 30 min, but was much lower at 60 min, suggesting that EGFR was transported to recycling endosomes. Similarly, EGF has been shown to display a lower signal at 60 min after treatment with recycling ligands, likely reflecting their combined trafficking at this high treatment concentration with the ligand^31^. Despite weak endosomal EGFR signal, the pAkt signal was continuously activated at least until 60 min. In TGFα-stimulated cells, significant pAkt signal was observed for as long as 180 min. This is probably due to continuous activation of the recycled receptors. In the case of EREG- and EPGN-stimulated cells, however, no significant change with regard to ligand stimulation were detected for the GFP-EGFR signal. This is likely due to low levels of phosphorylation/internalization of EGFR by EREG and EPGN, as compared with EGF^14^. Nevertheless, in the EREG-, but not the EPGN-stimulated cells, ligand addition rapidly increased pAkt signal, which is probably attributed to a distinct usage of signal transducers by EREG and EPGN^14^.

**Figure 1.**
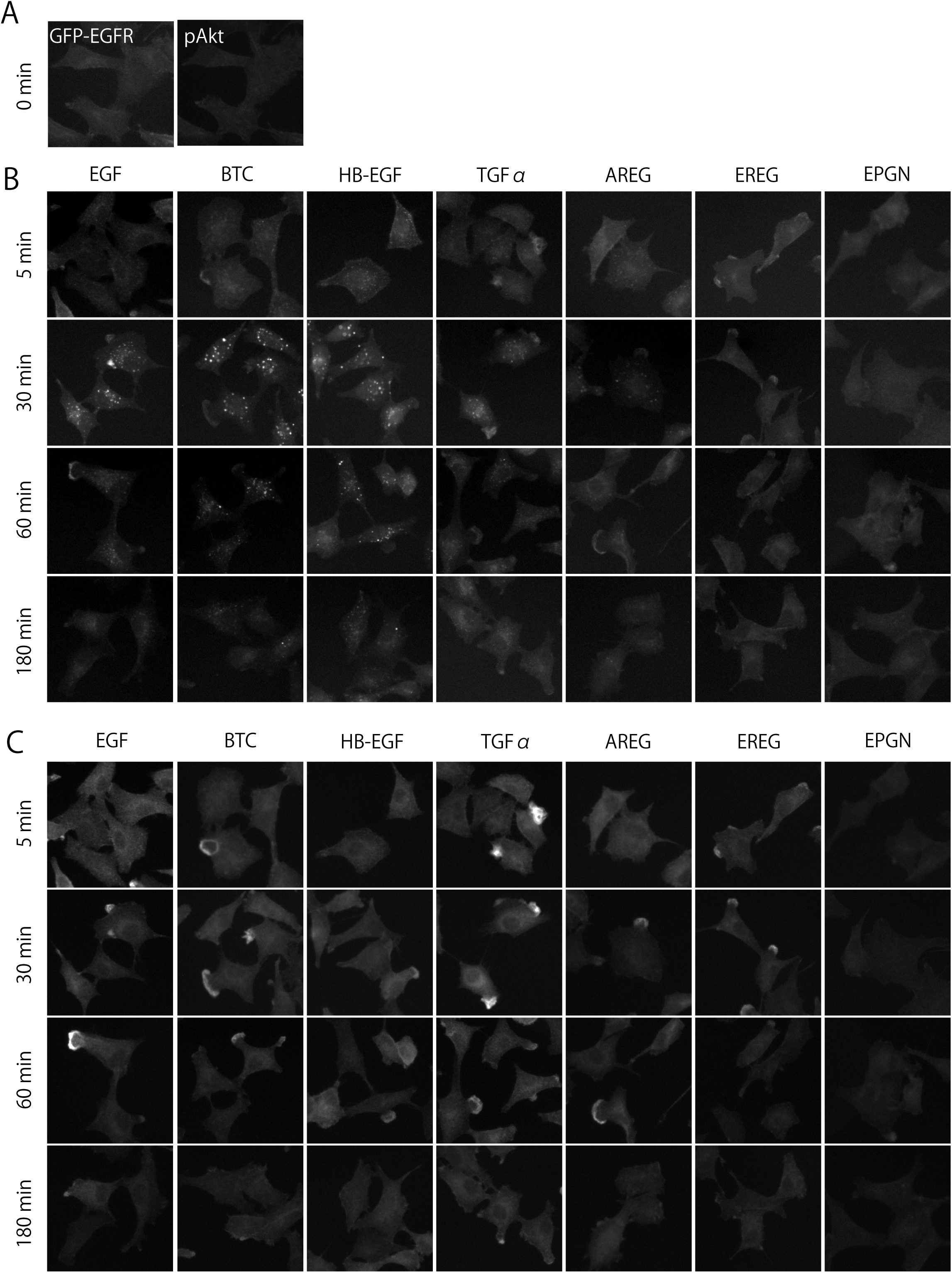
Representative images of EGFR ligand-stimulated cells. A549 cells expressing GFP-EGFR were serum-starved for 6 h and stimulated with EGFR ligands. Cells were fixed at the indicated time intervals following ligand stimulation, and processed for immunofluorescence to visualize GFP-EGFR (A) or phosphorylated Akt (pAkt, B).

Thus, differential cellular responses that were induced by different EGFR ligands were recognized at the level of microscopic images. Based on the above findings, EGFR ligands may be more accurately re-classified into three groups: degradative ligands, including HB-EGF and BTC, recycling ligands, including TGFα and AREG, and “recycling and weakly activating ligands”, including EREG and EPGN. The third group may be further discriminated by the level of signal transduction. EGF showed an intermediate (mixed) phenotype both of degradative and recycling ligands.

### Differential cellular responses triggered by different EGFR ligands can be quantitatively detected by image-based phenotypic profiling/HCS

Next, we attempted to quantitatively evaluate the cellular responses using image analysis as previously described^17^. Image features represented as integrated signal intensity, signal co-localization, and signal ratio were measured for each subcellular ROI (see Supplementary Table S1). The regions ‘Nucleus’ and ‘Cell’ were segmented according to the Hoechst staining and the GFP-EGFR signal, respectively. The ‘Cytosol’ region was defined by subtracting ‘Nucleus’ from ‘Cell’. Furthermore, the ‘Perinucleus’ and ‘Plasma membrane’ regions were obtained by the expansion of the region ‘Nucleus’ and the shrinking of the region ‘Cell’, respectively.

Representative image features are shown in Fig. 2A. Considering that Akt is phosphorylated by PDK1 at the intracellular side of the plasma membrane^33^, the integrated intensity of pAkt at plasma membrane (PM-pAkt) was consistent with what was expected (Fig. 2A, upper panels). Stimulation with any ligand (except EPGN) significantly increased PM-pAkt, with a peak intensity observed at 5 min. Following the peak, however, PM-pAkt behaved rather differently for the various ligands. In the HB-EGF- and BTC-stimulated cells, PM-pAkt decreased at 180 min, whereas, in the EGF- and TGFα-stimulated cells, PM-pAkt was continuously activated between 30 min and 180 min. Since stimulation with EGF at 100 ng/ml promotes both the degradation and recycling of EGFR^31^, the sustained activation of Akt may be attributed to the recycled EGFR. Of these ligands, stimulation with EPGN did not result in any substantial activation of PM-pAkt, consistent with the result presented in Fig. 1.

**Figure 2.**
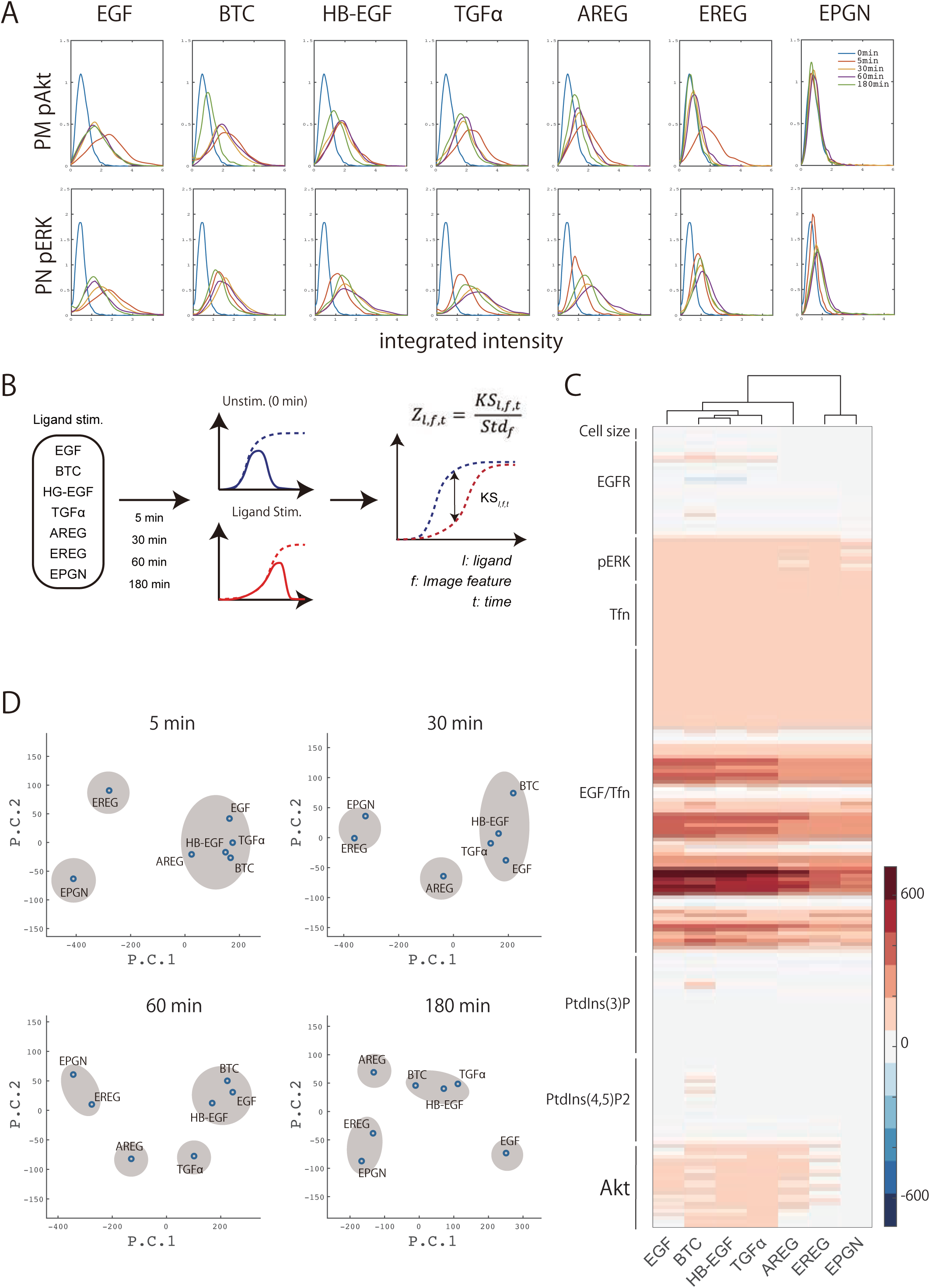
Quantitative cellular responses of EGFR ligand-stimulated cells. Ligand-stimulated cells were visualized for pAkt and pERK expression using immunofluorescence and quantitatively evaluated with image analysis. The integrated intensity of pAkt or pERK was extracted around the plasma membrane or perinuclear region, respectively (indicated as PM-pAkt and PN-pERK). The probability density function for each sample was evaluated using kernel density estimation (A). A two-sample Kolmogorov-Smirnov (KS) test between unstimulated (0 min) and ligand-stimulated cells was calculated for all the image features obtained from the image analysis (B). KS statistics were standardized (Z-score), and displayed as a heatmap (C). The Z-scores were subsequently reduced to account for redundancy using principal component analysis, and then processed for hierarchical cluster analysis to group the EGFR ligands (D).

The activation of the MAPK pathway, another major EGFR-initiated signaling pathway, can be monitored by observing the levels of pERK^34^. Since pERK translocates from the cytosol to the nucleus, pERK was measured for the perinuclear region (PN-pERK; Fig. 2A, bottom panels). PN-pERK increased gradually, reaching a peak intensity at 30–60 min after treatment with some of the degradative ligands (HB-EGF and BTC) or at 60 min after treatment with the recycling ligands (TGFα, AREG, and EREG). EGF induced a rapid increase in PN-pERK, which peaked at 5 min. EPGN, which induced no obvious activation of PM-Akt by phosphorylation, increased PN-pERK levels slightly, likely as a result of independent activation of receptor internalization^35,36^.

**Figure 3.**
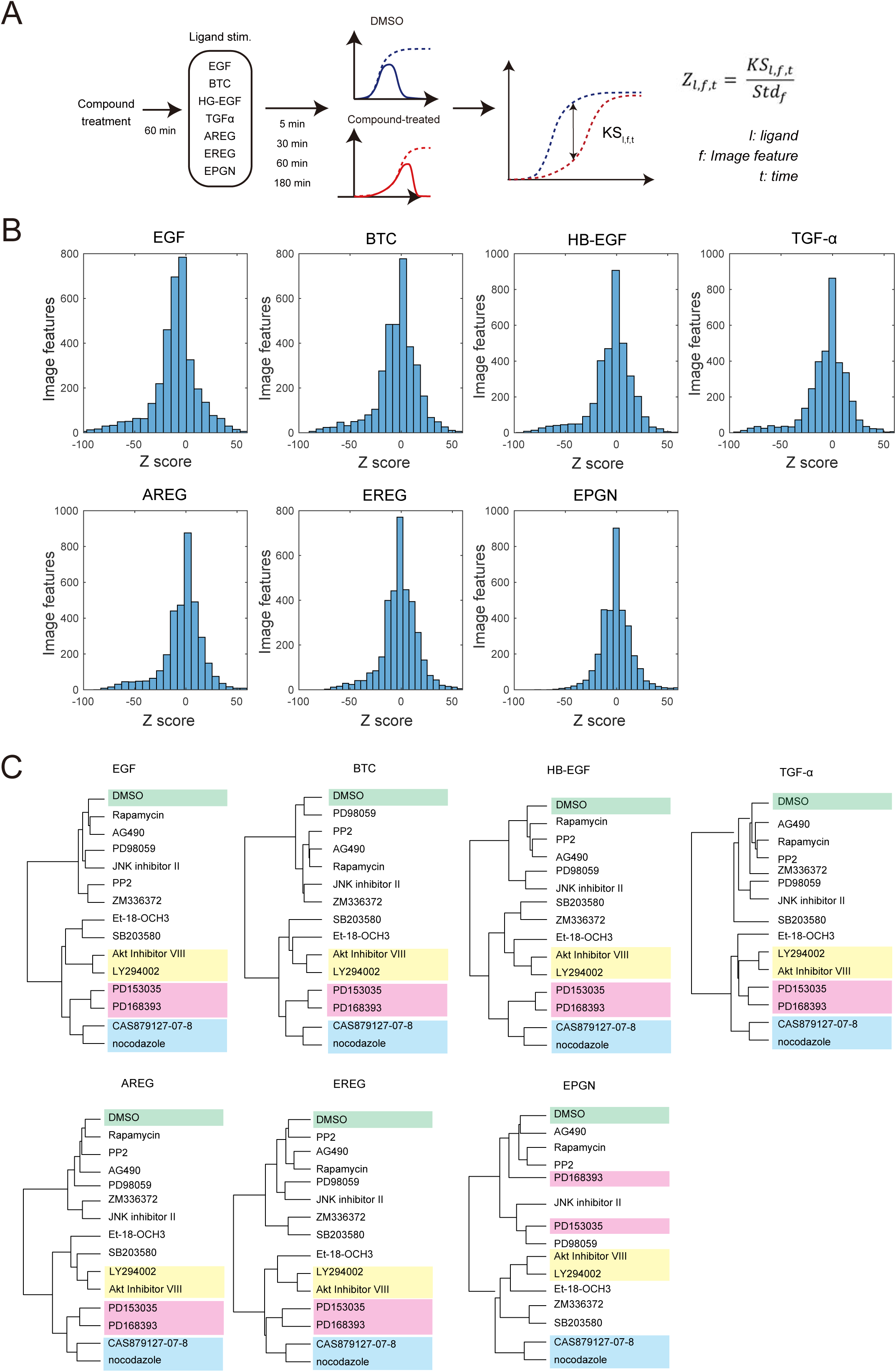
Image-based profiling of compounds under differential stimulation. Cells were serum-starved for 5 h, and incubated with inhibitors for 1 h. Cells were stimulated with ligands for the indicated time periods (5, 30, 60, 180 min), and subsequently processed for immunofluorescence. A two-sample KS tests was performed for DMSO- and inhibitor-treated cells for each ligand, and used to calculate selected Z-scores as described in the materials and methods (A). To evaluate the overall effect of the inhibitors, the Z-score distribution for each ligand is shown (B). To reduce redundancy of the selected Z-scores, principal component analysis (PCA) was performed. Six principal components (PCs), which account for more than 95% of the variance, were used for hierarchical cluster analysis (C).

Image features of ligand-stimulated cells were compared with those of unstimulated cells using a modified two-sample KS test^6,17^ (Fig. 2B). The KS statistics were standardized (Z-scores), and an average of two independent experiments was obtained (mean Z-scores). A heatmap of the mean Z-scores is shown in Fig. 2C. Various degrees of both similar and different cellular responses were induced by each EGFR ligand. The bars alongside the panel indicate the signaling molecules with which the image features are associated. Next, we attempted to classify these ligands based on their cellular responses by using two principal components and plotting each ligand in two-dimensional space (Fig. 2D). EREG and EPGN were separated from other ligands and from each other at 5 min, corresponding to their ability to slightly induce EGFR trafficking and differential Akt activation (see Fig. 1 and Fig. 2A). Since TGFα and AREG induced significant EGFR trafficking and signal activation, these two ligands could not be discriminated from the degradative ligands at 5 min. However, after 30–60 min, these two ligands were separated from the degradative ligands, likely caused by TGFα- and AREG-mediated induction of EGFR recycling. Two degradative ligands (HB-EGF, BTC) and EGF were grouped together until 60 min, but EGF was separated from the other two (HB-EGF and BTC) at 180 min. This may be a result of EGF-bound EGFR signaling through both the degradation and recycling pathways^31^.

The results presented in Fig. 2 collectively indicate that image-based phenotypic profiling/HCS can quantitatively differentiate cellular responses induced by different EGFR ligands.

### Image-based phenotypic profiling of compounds under various ligand stimulation

Next, we attempted image-based phenotypic profiling of compounds under various ligand stimulation. The compounds examined were inhibitors associated with the EGFR pathway: Akt inhibitor VIII for Akt, PD153035, PD168393, and CAS879127-07-8 for EGFR, Et-18-OCH3 for PI-PLC, JNK inhibitor II for JNK, LY294002 for PI3K, PD98059 for MEK, PP2 for Src, Rapamycin for mTOR, SB203580 for p38MAPK, AG490 for JAK2, ZM336372 for c-Raf, and nocodazole for microtubules. Of these 14 compounds, CAS879127-07-8 inhibits microtubule polymerization as well as EGFR kinase activity^17^. The cells were treated with each inhibitor for 1 h, stimulated with each EGFR ligand, and processed for image-based phenotypic profiling as described above. The combination of each inhibitor (with DMSO as a control treatment) and ligand yielded 105 samples (15 × 7 = 105). All the samples were processed for staining in triplicate at four time points (105 × 3 × 4 = 1260 samples).

The effects of each inhibitor on PM-pAkt and PN-pERK levels are presented in Supplementary Fig. S1 and S2, respectively. Similar patterns of a given compound’s effects on image features were observed irrespective of the use of different ligands. To quantitatively evaluate the inhibitor’s effects, image features from control (DMSO)- and inhibitor-treated cells at the same time point were statistically compared using a two-sample KS test (Fig. 3A). As described above, KS statistics were standardized and averaged using two independent experiments (mean Z-scores). The distribution of the mean Z-scores for each ligand-stimulated sample is shown in Fig. 3B. Of these, the Z-scores in the EPGN-stimulated cells were narrow in range, suggesting that the inhibitor’s effects were limited. This is likely related to the low activation by EPGN (see Supplementary Fig. S1 and S2). In the other ligand-stimulated cells, the distribution of the selected Z-scores was broad and within similar ranges, suggesting that the amplitude of inhibitor effects is likely comparable.

Next, to reduce the redundancy of image features, PCA was carried out using the selected Z-scores. All 14 compounds treated with each ligand were classified by hierarchical cluster analysis using 14 PCs (Fig. 3C). Interestingly, under any ligand stimulation except for EPGN, nearly half of the compounds were classified in the same category (e.g., a cluster of PI3K-Akt inhibitors (LY294002 and Akt inhibitor VIII), a cluster of EGFR inhibitors (PD153035 and PD168393), and a cluster of microtubule inhibitors (nocodazole and CAS879127-07-8)). Thus, the profiling of compounds known to inhibit EGFR signaling was not greatly affected by differential EGFR ligand stimulation.

On the other hand, hierarchical clustering of compounds under the stimulation with EPGN resulted in only two clusters, PI3K-Akt inhibitors and microtubule inhibitors. Two EGFR inhibitors, PD153035 and PD168393, did not form a cluster. Since EPGN induces weak cellular responses, as shown in Fig. 1 and 2, these two EGFR inhibitors may have exerted limited effects on EGFR itself, but displayed unexpected secondary effects.

### Ligand differences do not affect compound profiling

Finally, compound profiling results were quantitatively compared among the different ligand treatments. The cosine distance between the control (DMSO) and inhibitor treatment was calculated for each inhibitor and ligand stimulation, and used as an index of similarity (or difference) in the inhibitor’s effect. The distances were plotted for each pair of ligands (21 panels from 7 EGFR ligands, Fig. 4). Correlation coefficients between the two ligands were calculated and are shown in each plot. For most ligand combinations, the correlation coefficients were greater than 0.9. This indicates that the observed effects of a given compound when stimulated with different ligands were not significantly different from each other.

**Figure 4.**
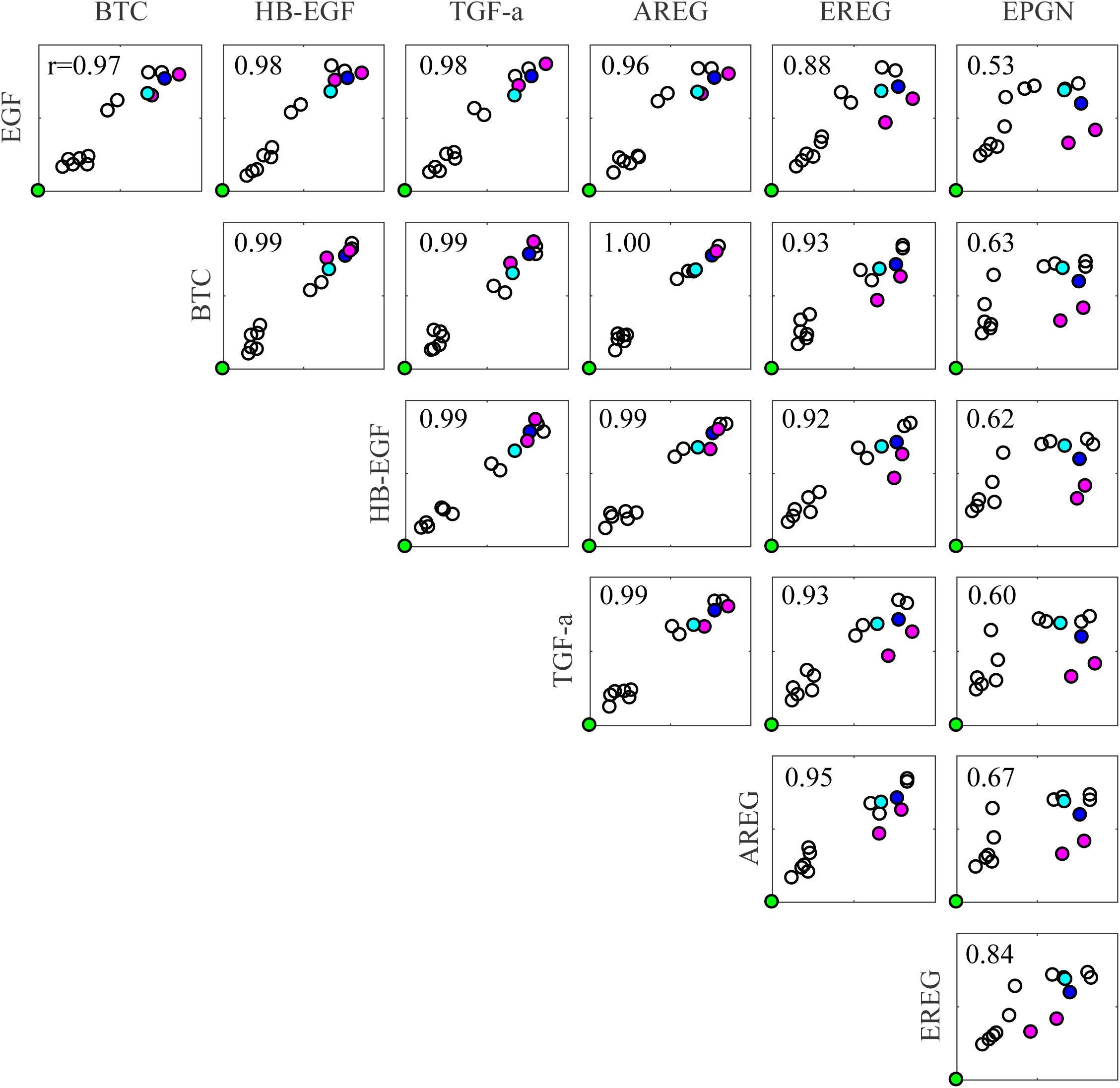
Similarities and differences of compound effect among differentially stimulated samples. Cosine distances of six PCs between the control (DMSO)- and inhibitor-treated samples were calculated and plotted for each ligand. The correlation coefficients (r) for each ligand combination are displayed in each plot. Green represents the control (DMSO) treatment, magenta indicates the two EGFR inhibitors (PD153035 and PD168393), blue corresponds to CAS879127-07-8, and cyan represents nocodazole.

The combination of a given ligand with EPGN, however, displayed low correlation coefficients (∼0.6). In all of the scatter plots involving EPGN, two common outliers were observed, corresponding to EGFR inhibitors (PD153035 and PD168393, colored by magenta in Fig. 4). Their emergence as outliers may be related to EPGN exhibiting low levels of EGFR activation, so the effects of EGFR inhibitors on EGFR were rather limited. CAS879127-07-8 (blue in Fig. 4), another EGFR inhibitor, displayed a similar effect, even in EPGN-stimulated cells. This is likely due to its inhibitory effect on microtubules (cyan). The result is consistent with the hierarchical clustering of compounds (Fig. 3C).

## Discussion

In image-based phenotypic profiling/HCS, many image features are utilized to evaluate cellular phenotypes. Using this approach, we previously revealed that a highly specific kinase inhibitor of EGFR, CAS879127-07-8S, also functions as a microtubule inhibitor^17^. This is an example demonstrating that this method can become a powerful tool for discovering new drug action mechanisms^6,10,11^.

However, there is concern that experimental conditions can affect the result of compound profiling by image-based phenotypic profiling/HCS. In the present study, we investigated this possibility using seven different EGFR ligands^13,16^. These ligands bind to the same receptor (EGFR) but induce differential signal transduction and cellular responses, which can be readily detected with image-based profiling. For each ligand, 14 compounds, all known to inhibit various EGFR signaling pathways, were evaluated with image-based profiling, as previously described^17^. Interestingly, for all the ligands except EPGN, nearly half of the compounds were classified into groups that were consistent with their established target molecules. They correspond to a cluster of EGFR inhibitors, a cluster of PI3K-Akt inhibitors, and a cluster of microtubule inhibitors. Furthermore, the correlation coefficient indicating the compounds’ effects for a combination of two ligands (excluding EPGN) was very high (>0.9). Thus, image-based compound profiling is quite robust, even with some variance in experimental conditions.

It is interesting that the use of different ligands did not greatly affect the compound profiling. In particular, PI3K-Akt and microtubule inhibitors were distinguished from the others irrespective of the ligand choice (including EPGN). One possibility is that the characteristics of these two clusters of inhibitors may be determined by image features that were not related to EGFR stimulation. For example, since PtdIns(3)P can easily be detected prior to ligand stimulation and was reduced by the PI3K inhibitor treatment, the cellular effects of the PI3K inhibitor were discernible in the absence of EGFR activation. Similarly, the basal level of PM-pAkt at 0 min was sufficient to be detected and was significantly decreased by Akt inhibitor VIII and LY294002. Thus, basal PM-pAkt expression may contribute to the clustering of Akt inhibitors, even without EGFR activation. Microtubules are involved in transferrin trafficking^32,37^. Therefore, image features associated with transferrin are sufficient for the discrimination of microtubule inhibitors from other inhibitors in the absence of EGFR activation.

In the case of EPGN-stimulated cells, two EGFR inhibitors were not classified into a single cluster. Since EPGN induces both EGFR endocytosis and its downstream activation^16,38^, the effects of EGFR inhibitors on the EPGN-EGFR pathway were most likely limited, and their potential secondary effects may have caused their separation from each other. Conversely, with the exception of EPGN treatment, two EGFR inhibitors were classified into the same cluster, likely reflecting their authentic effects on EGFR activation.

This study demonstrated that cell stimulation with various EGFR ligands can be detected by profiling image features and that the profiling of various signaling inhibitors was not largely affected by the specific ligand. While more research is necessary to further validate the robustness of HCS in different conditions, this study provides a valid basis for its extended application in the future.

## Acknowledgments

The authors thank A. Inagaki for technical assistance and discussions. This work was supported by JSPS KAKENHI grant number JP17K07347 and the Takeda Science Foundation.

